# “A Low-Cost Microfluidics System for Light Microscopy Experiments”

**DOI:** 10.1101/2024.04.29.591694

**Authors:** Logan M. Harrison, Benjamin M. Brisard, Kylie D. Cashwell, Aaron L. Mulkey, Cameron A. Schmidt

**Affiliations:** Department of Biology, East Carolina University, Greenville NC

## Abstract

Microfluidics devices are powerful tools for studying dynamic processes in live cells, especially when used in conjunction with light microscopy. There are many applications of microfluidics devices including recording dynamic cellular responses to small molecules or other chemical conditions in perfused media, monitoring cell migration in constrained spaces, or collecting media perfusate for the study of secreted compounds in response to experimental inputs/manipulations. Here we describe a configurable low-cost (channel-based) microfluidics platform for live-cell microscopy, intended to be useful for experiments that require more precision/flexibility than simple rubber spacers, but less precision than molded elastomer-based platforms. The materials are widely commercially available, low-cost, and device assembly takes only minutes.

## Introduction

Microfluidics devices are broadly used to facilitate flexible experimentation in quantitative cell biology[1]. Channel based microfluidics devices mediate the flow of fluids through a channel or system of channels, and this simple approach can be used for many research applications including systematically changing perfusate media composition while monitoring biological functions, developing chemical gradients or fluid mixing regimes, exposing cells to sheer forces, etc. A few common materials are typically used in microfluidic device fabrication including-elastomers (e.g. polydimethylsiloxane (PDMS)), thermoplastics (e.g. polyethylene terephthalate (PET), and paper[2].

Channel-based microfluidics devices are typically constructed either by molding a channel into a material (commonly PDMS) and chemically bonding the material to a cover, or by cutting a channel shape out of a spacer material and placing the spacer between two pieces of optically clear plastic or glass. In both cases, fluid in-flow and out-flow channels are added, and fluid is perfused using a programmable syringe or peristaltic pump. The degree of sophistication and precision of channel-based microfluidics devices varies widely among commercial platforms, from simple devices consisting of rubber spacers applied to a microscope slide, to sophisticated molded devices with light actuated valves[3]. Simple rubber spacers can be problematic when optical resolution in the axial plane is important for the experiment, because they are relatively thick. Additionally, commercially available designs are limited. More sophisticated PDMS chips are much more precisely shaped at the microscopic scale and are more amenable to flexible design but are costly, time consuming to fabricate, and require specialized equipment (e.g., plasma cleaning, soft lithography, etc.).

In this report, we describe a low-cost channel-based microfluidic device fabrication system aimed at providing flexible devices with moderate precision for live cell microscopy applications that demand more than simple rubber spacers, but less than highly sophisticated elastomer-based devices. We assess three use cases: 1.) modeling fluid mixing rates in the channels, 2.) performing stop-flow ‘micro-insemination’ experiments with live mammalian spermatozoa, and 3.) performing time-series video microscopy on adherent mouse myoblasts (C2C12 cells).

## Materials and Methods

### Chemicals and Reagents

Chemicals and reagents were obtained from Sigma-Aldrich (St Louis, MO). Several specific media were used in this study. Human tubal fluid minimal media (HTF-min) consisted of (mM): NaCl (120), potassium phosphate monobasic (0.35), magnesium sulfate heptahydrate (0.4), potassium chloride (4.7), Bovine serum albumin (BSA; 5 g/L), pH 7.4 at 37°C. Sperm isolation media consisted of HTF-min supplemented with EGTA (ethylene glycolbis(β-aminoethyl ether)-N,N,N′,N′-tetraacetic acid) (1). Capacitation media consisted of HTF-min supplemented with sodium bicarbonate (25) and calcium chloride (2.5). Complete media consisted of capacitation media supplemented with sodium pyruvate (0.33), sodium L-lactate (21), and D-glucose (2.78)[4].

### Modeling Fluid Mixing

To model fluid mixing rates, we tested two separate mixing experiments-1.) using resazurin dye measured spectrophotometrically at 610 nm, compared to a dilution standard curve, and 2.) using glucose measured via a glucose-hexokinase enzyme coupled reaction scheme, also calibrated to a dilution standard curve (ThermoFisher, Infinity Glucose Hexokinase Reagent). The initial perfusate did not contain either reagent, and was switched to a new source containing either resazurin or glucose after the microfluidic device was fully filled (t=0). In both experiments, the perfusate was collected in the wells of a 96-well microtiter plate with 50 μL of perfusate in each well under a constant flow rate of 25 μL/minute. Measurements were made using a microtiter plate reader (ID3, Molecular Devices, San Jose, CA).

### Animals

Adult male outbred CD-1 mice, aged 16–24 weeks old, were obtained from Charles River Laboratories (Raleigh, NC, USA). All work adhered to the guidelines outlined in the National Research Council Guide for the Care and Use of Laboratory Animals and was approved by the Institutional Animal Care and Use Committee of East Carolina University (approval A3469-01). Mice had free access to water and food, were maintained on a 12 hr light/dark cycle and were humanely euthanized by CO_2_ asphyxiation followed by thoracotomy.

### Imaging Live Mouse Epididymal Spermatozoa

Testes with intact epididymides were isolated in phosphate buffered saline (PBS) at 37°C. Cauda epididymides were transferred to isolation media where they were cut open. Following a brief swim-out period (∼15 mins at 37°C), tissue was removed, and sperm were isolated by centrifugation at 800 x g for 2 mins followed by at least one wash step. Cell counts were determined using a hemocytometer after dilution in water. Microfluidics devices were fabricated and pre-equilibrated in HTF media for 15 mins on a thermal stage (Linkam Scientific Instruments, Salfords, Redhill, UK). Spermatozoa were injected into the microfluidic device antechamber at a flow rate of 25 µL/min. Flow was then stopped and spermatozoa were monitored in the channel space for 10 mins to assess cell migration at various positions. Cells were imaged using a Zeiss Axiovert.A1 with a 10X magnification negative phase contrast achromat objective (NA = 0.25) (Carl Zeiss AG, Jena, Germany). Zeiss Zen Blue software with a time-controller add-on was used for acquisition.

### Imaging Mouse C2C12 Myoblast Cells

C2C12 cells were purchased from ATCC (CRL1772; Manassas, VA, USA) and subcultured in Dulbecco’s Modified Eagle Medium with 25 mM D-glucose, 4 mM L-glutamine, and 10% v/v fetal bovine serum. Subcultured cells at passage six were trypsinized from a T-75 culture flask centrifuged at 400xg for 5 mins to remove trypsin solution and injected into a microfluidics chamber for live imaging at a density of 10^3^ cells/cm^2^. Cells were imaged using a Zeiss Axiovert.A1 with a 10X magnification negative phase contrast achromat objective (NA = 0.25).

## Results

### Design

Several alternative designs and methods of construction were attempted before arriving at the final version presented in this report. Primary challenges included: 1.) obtaining a water-tight seal between the cover slides and spacer, 2.) preventing accumulation of air bubbles during perfusion, and 3.) identifying appropriate slide material to facilitate imaging with phase contrast microscopy-which is extremely sensitive to slight variations in the refractive index of the materials. The optimized (final) version of the device consists of a spacer made of double-sided adhesive with a central cutout that can be flexibly designed using a digital die-cutting machine (Cricut Joy® and Cricut Design Space® software) (**Figure 1**). The spacer and cutout design provides a small perfusion channel when placed between two slides. The top slide in the constructed device was modified to include two tubing adaptors for placement of inflow and outflow channels. Perfusion through the device can be achieved using any appropriate syringe pump (non-peristaltic in this report) and an effluent collection chamber.

**Figure 1:**
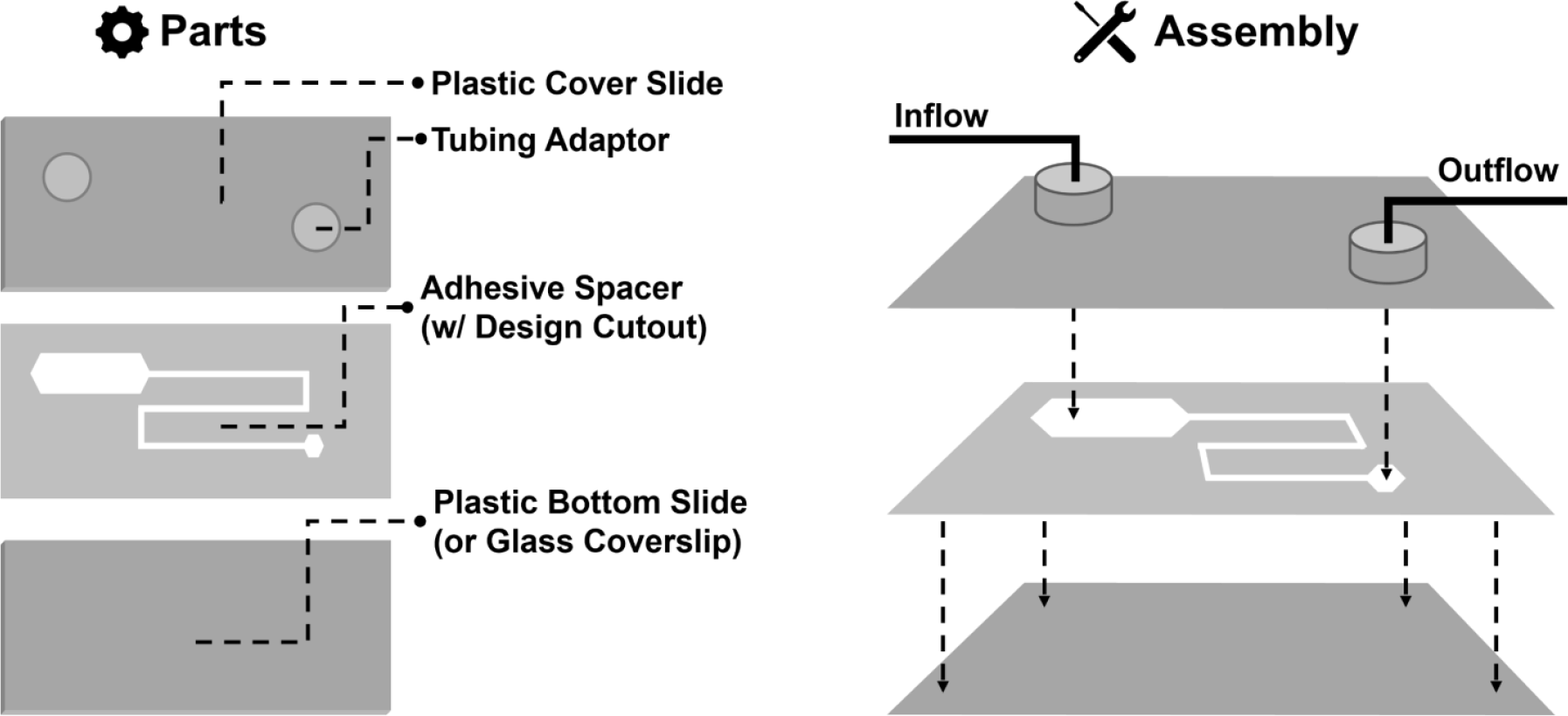
Device Design Schematic.

### Fabrication

The cover slides consist of two pieces of optically clear polyethylene terephthalate (PET) plastic, cut into the shape of microscope slides by scoring/breaking with a hobby knife. A spacer is designed according to user needs and cut using a hobby die-cutting machine. The smallest distance between two parallel lines cleanly cut by the machine was determined to be approximately 1 mm, enabling the production of small channels. The spacer is made from a sheet of double-sided adhesive. The thickness of the adhesive was measured with a micro caliper and determined to be approximately 0.08 mm thick. The spacer, once cut, was then ‘seeded’ with a small needle or pointed tool. The seeding process involves removing the unwanted adhesive material and backing from the inner portion of the spacer. Once seeded, the spacer was then transferred to the bottom plastic slide.

The top plastic slide was modified to accommodate the input and output flow connections. Holes for the flow connections were drilled using a battery-operated rotary tool with a 1 mm diameter diamond coated drill bit. A small drop of water was placed over each drilling site to prevent the formation of plastic dust. Once drilled, the slides were then placed together and sealed using a heavy flat object. It was necessary to apply a large amount of pressure to ensure a strong seal and prevent fluid from escaping the desired flow path through the device. Notably, during device operation, high perfusion pressures or poor initial seal resulted in small amounts of fluid escaping from the sides of the device in some rare cases. The input/output flow connections were constructed using PE-50 tubing inserted into adhesive silicone tubing adaptors (Grace Biolabs). To pull the tubing through the adaptors, a 21 ½ gauge blunt tipped needle was inserted into the adapter, and the tubing was fit over the needle. The tubing was then pulled through the adapter until approximately 0.5-1.0 mm of tubing protruded from the side with the adhesive. The tubing adaptors with the tubes attached could then be applied to the holes drilled in the top layer of PET plastic in the device, using the protruding 1mm of tube to aid in alignment. The complete construction of the devices is illustrated in (**Figure 2)**.

**Figure 2:**
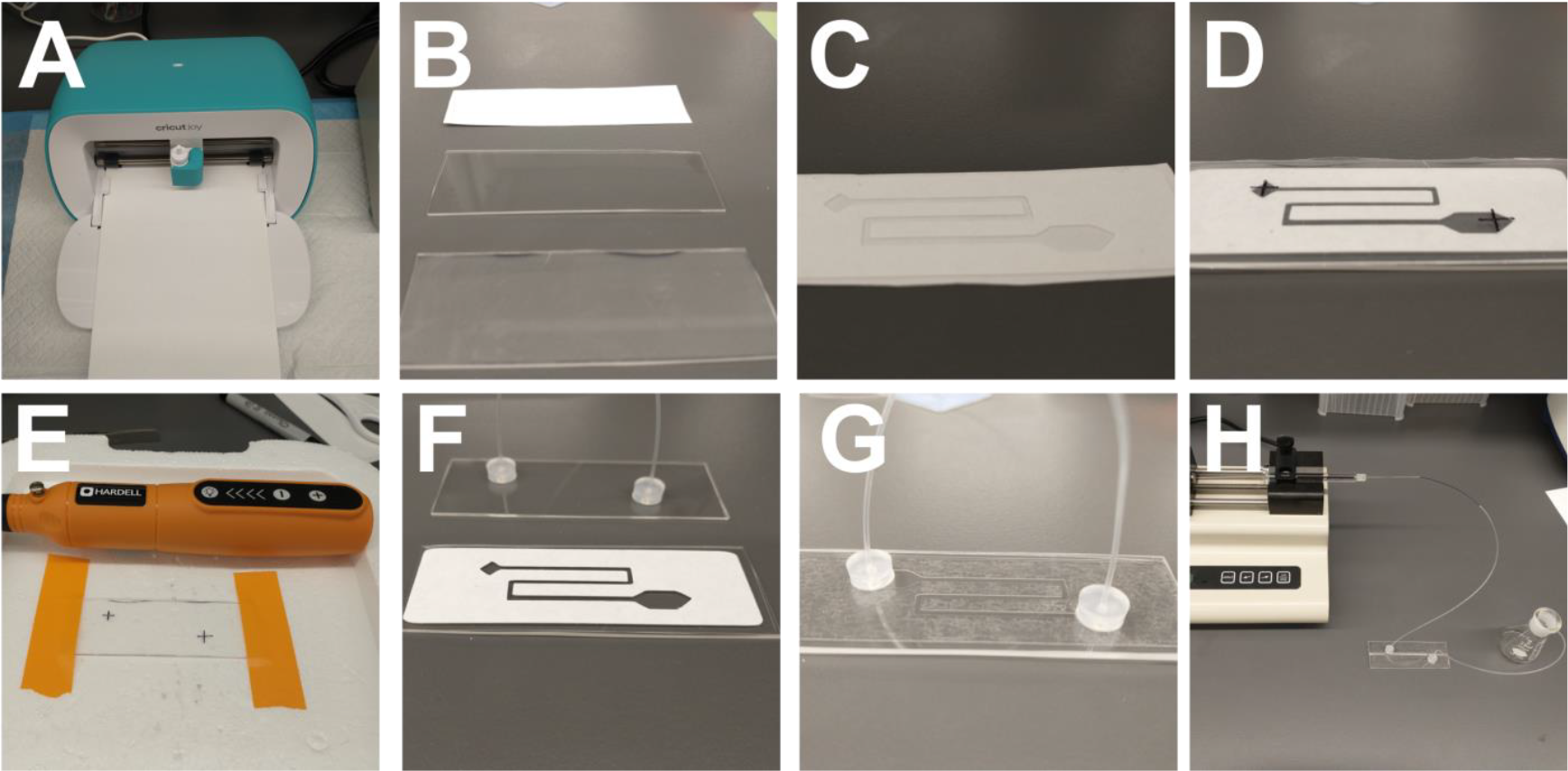
Device fabrication steps. (A) Design/cut the spacer channel shapes using the Cricut® die-cutting device with the Design Space® software. (B) Cut the cover slides and spacer to an appropriate size. (C) ‘Seed’ the spacer and remove backing. (D) Apply the spacer to the bottom slide and align the top slide to mark for drilling holes. (E) Drill the holes in the top slide. (F) Attached the tubes/tubing adaptors to the top slide. (G) Assemble the device. (H) Connect to in-flow and out-flow tubing and perfuse at an appropriate flow rate (determined for each application).

### Components and Cost

Specific components, costs, and sources are listed in (**table 2**). Because the materials are purchased in bulk (relative to the material required for a single device), the per-device cost is less than one dollar. The highest unit cost materials are the PE-50 tubing and the syringe pump. Notably, open-source plans are widely available for building low-cost syringe pumps if necessary[5].

**Table 1:**
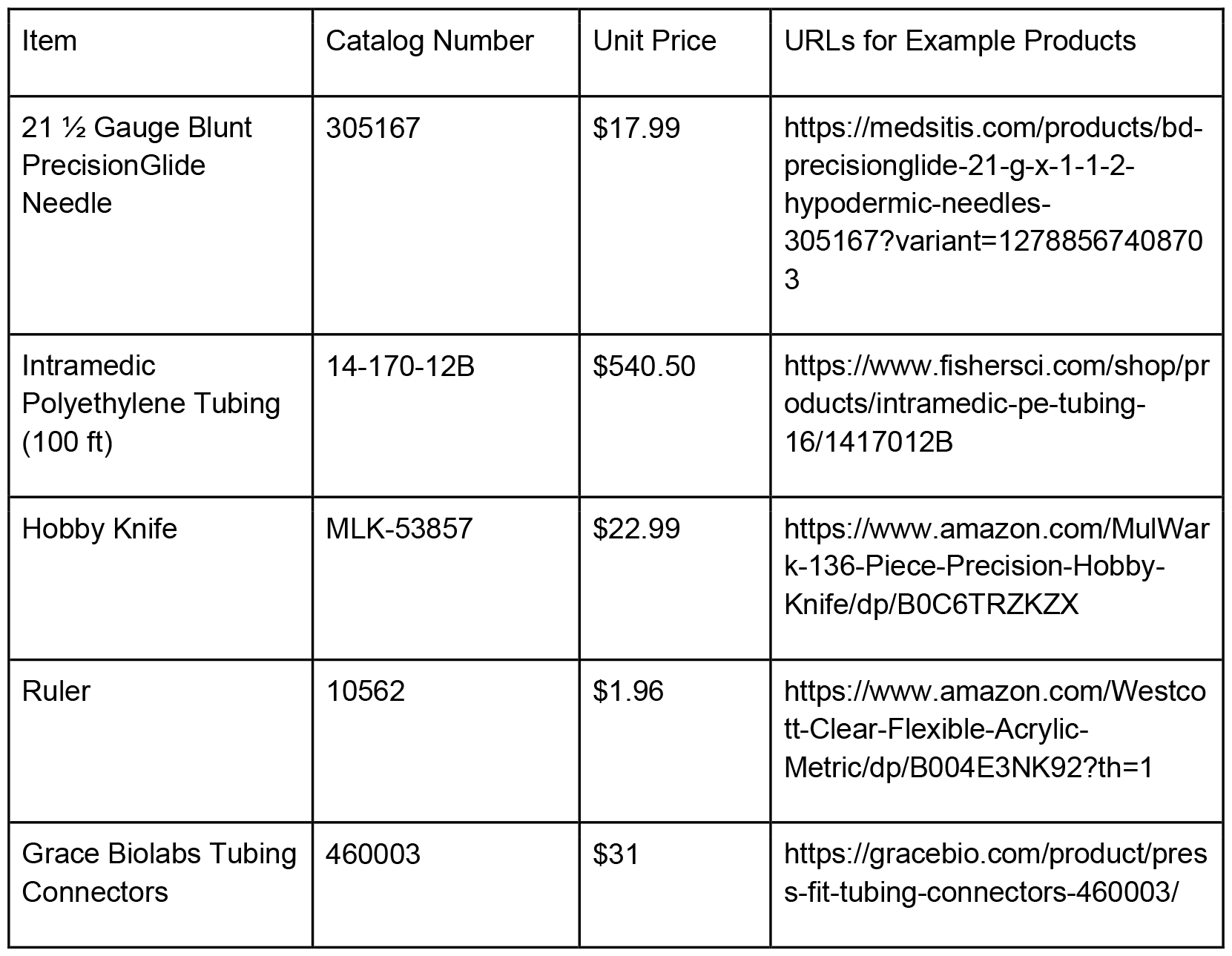
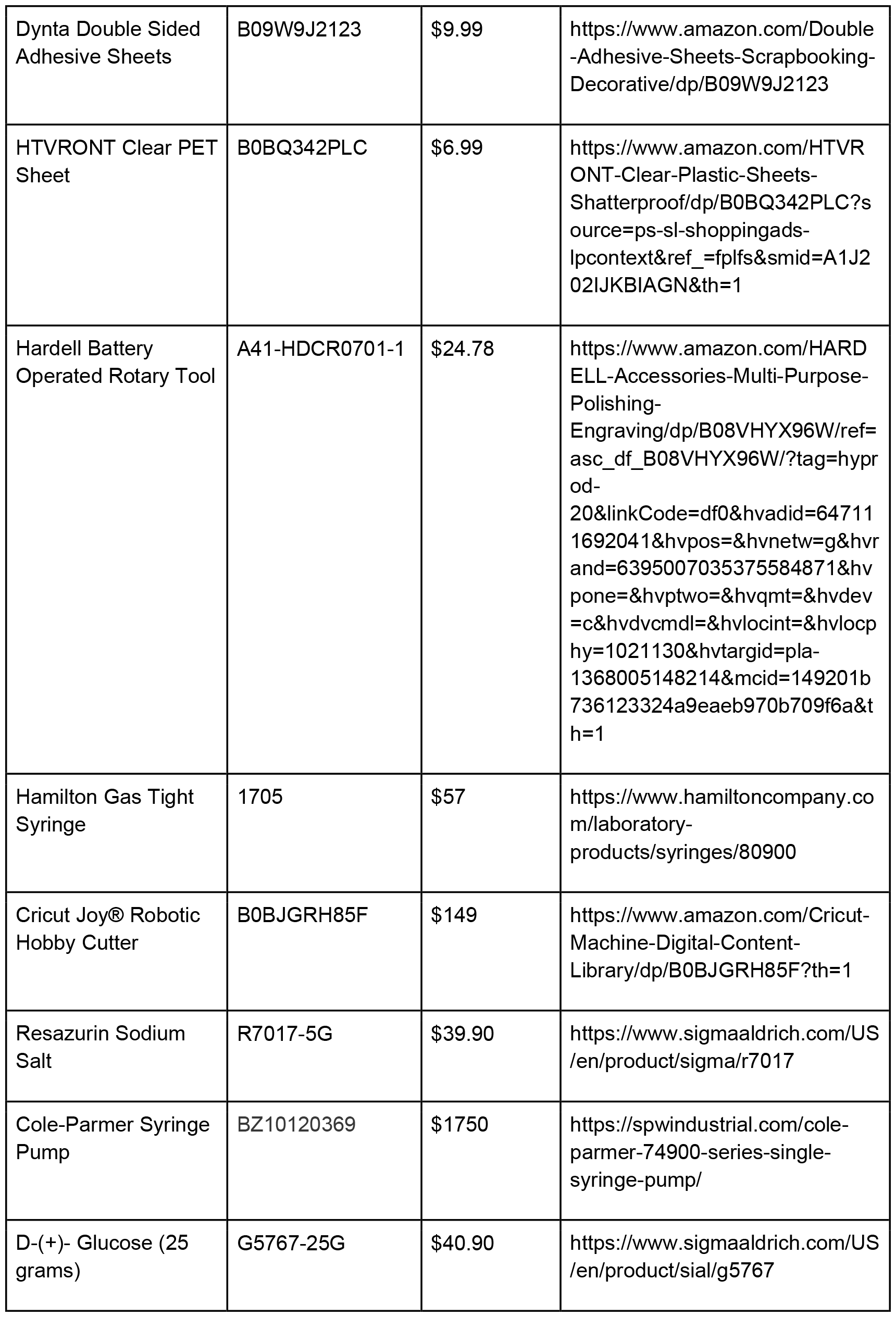
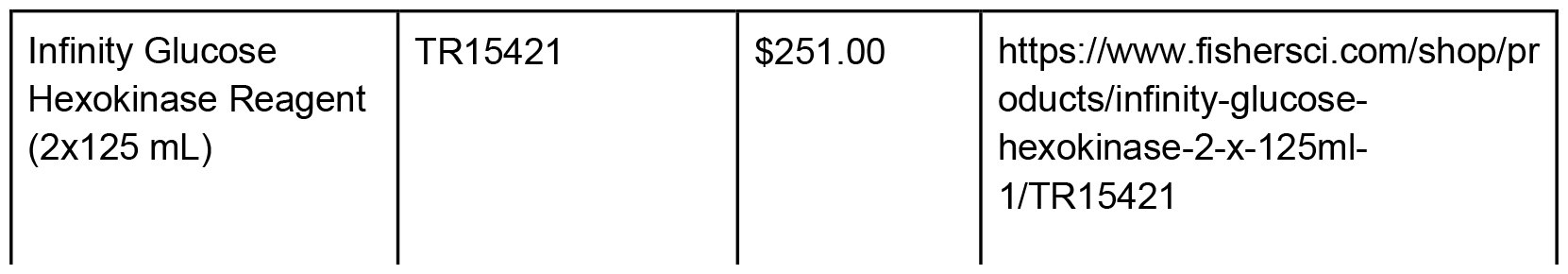
Summary of Device Components and Cost.

### Modeling Fluid Exchange Rates

A potential use of these microfluidic devices is to monitor live cells during a transition period in which the cells are exposed to changing reagents in the perfusate. For example, cells could be exposed to a glucose free media, and the perfusion source could then be switched to a media containing glucose allowing the concentration to increase in the middle chamber in accordance with the fluid properties and flow rate. In this design, the experimenter may want to model the mixing time of the reagent in the device so that the cells’ response to the reagent could be measured in a linear concentration range. With preliminary testing we were able to achieve a 10-minute window of (approximately) linear mixing, which was separately tested, and indicated using resazurin dye or perfusate glucose concentration (**Figure 3**). This approach could be further modified to suit specific purposes by adjusting the spacer cutout design, flow rate, or fluid properties. The testing method could be extended to any molecule of interest if it can be measured in the collected perfusate or optically in the channels of the device. Alternatively, the resazurin dye method could be used to measure/model the mixing dynamics if the molecule of interest cannot be measured directly.

**Figure 3:**
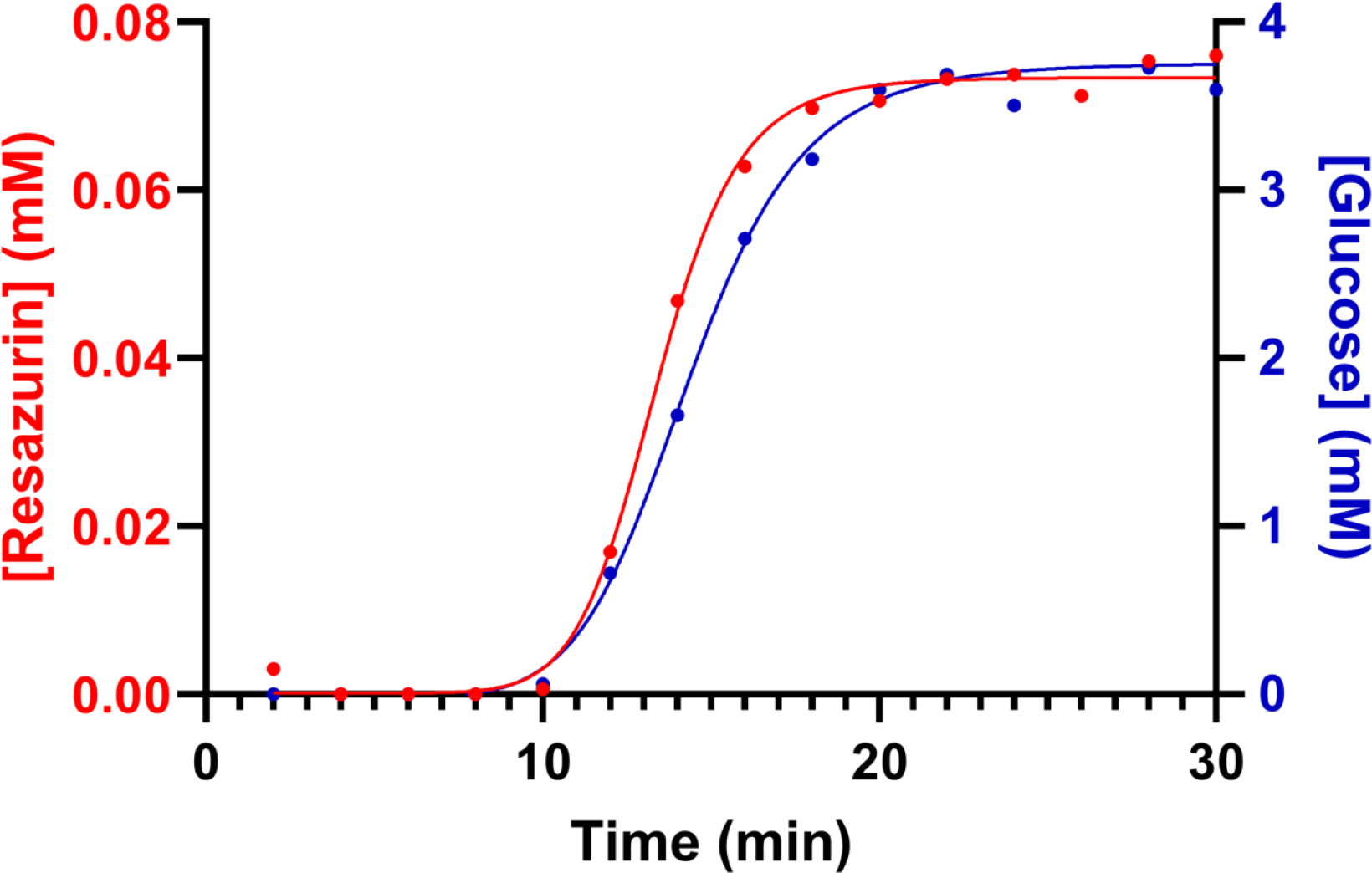
Modeling fluid mixing to identify a linear concentration range. (Left Y-axis) resazurin concentration measured over time in collected perfusate. (Right Y-axis) glucose concentration measured over time in perfusate. Flow rate = 25 µL/min.

### Stop-Flow Imaging of Live Mouse Spermatozoa

Spermatozoa exhibit motility patterns that are influenced by local fluid flows and physical fluid properties (e.g. viscosity and viscoelasticity). Additionally, motility analyses are typically performed in glass depth-chambered slides that do not involve navigation through constrained spaces. To study the utility of microfluidic devices for studying sperm motility dynamics, live mouse epididymal spermatozoa were isolated and injected into a device that was prefilled with human tubal fluid media. The device allows for ‘micro-insemination’ analysis in which a dense sample of spermatozoa are injected into an antechamber (**Supplementary Video S1**), the flow is then stopped, and the ability of the sperm cell population to explore user-defined complex spaces is then analyzed using video microscopy and accompanying computer vision strategies (**Supplementary Video S2**). In initial testing, a glass slide was used for the bottom of the device (the top must be plastic to facilitate drilling). Glass resulted in spermatozoa sticking to the surface, which only nominally mediated by changing the concentration of media bovine serum albumin. For this reason, plastic was used, which resulted in less cell adherence.

### Time-Lapse Imaging of Adherent C2C12 Myoblasts

To study adherent cells, mouse C2C12 myoblasts were injected into the device and allowed to adhere for 15 minutes in a 37C incubator. Cell movement was then monitored for one hour on an inverted microscope with a 37C heated stage while phase contrast images were recorded every 15 seconds (**Supplementary Video S3**). To assess whether low fluid flow rates would cause loss of cell adhesion, a continuous flow of subculture media (DMEM) was applied at a rate of 25 μL/min. No apparent loss of adhesion was observed on the plastic bottom slide. Note that the plastic slides are approximately 1 mm thick. If imaging with a low working distance objective is necessary, the bottom slide could be replaced with a coverslip of dimensions that suit covering the open channels within the spacer. If cell adhesion is determined to be poor on glass, a coating such poly-L-lysine could be applied as appropriate for user defined needs.

## Discussion

In this report, we describe a low-cost microfluidic device that can be used for various live-cell microscopy applications. These devices are directly suitable for applications involving suspended cells as well as adherent cells and can be adapted to virtually any imaging platform (e.g., using a glass coverslip as the bottom slide for imaging on low working distance objectives). The devices are also amenable to dynamic experiments involving fluid exchange for measuring cellular responses to media reagents over a defined concentration range.

The primary strengths of these devices include: 1) user defined channel design to suit a wide array of experimental needs, 2) low cost of device materials, and 3) relative ease of construction as well as quick construction time (only a few minutes). There are, however, several potential limitations of these devices. First, the spatial tolerance of the robotic die-cutting device is not precise at the microscopic scale and the narrowest channel width consistently achievable was ∼1 mm. Second, plastics and adhesives come into direct contact with the perfusion media during operation and depending on the composition/sources of these materials, there may be some toxicity to live cells (though we did not observe any apparent toxicity). Third, the device seals were only stable at low pressures, necessitating relatively low fluid flow rates. Though, this consideration largely depends on the double-sided adhesive material used in the spacer, and there may be more robust options that are commercially available.

## Author Contributions

LH, BB, KC, AM, and CAS performed device fabrication and testing experiments. LH and CAS wrote the manuscript. LH, BB, KC, AM, and CAS reviewed and edited the manuscript.

## Funding

This work was supported by the Eunice Kennedy Shriver National Institute of Child Health and Human Development (R01HD110170-01A1), as well as laboratory startup funding from the Thomas Harriot College of Arts and Sciences at East Carolina University and the East Carolina University Research and Economic Development Office.

## Disclosures

The authors declare no conflicts of interest.

## Acknowledgements

None

## Notes

### Competing Interest Statement

The authors have declared no competing interest.

