## Supplementary figures and images for "“A Low-Cost Microfluidics System for Light Microscopy Experiments”"

### Supplementary Video S1

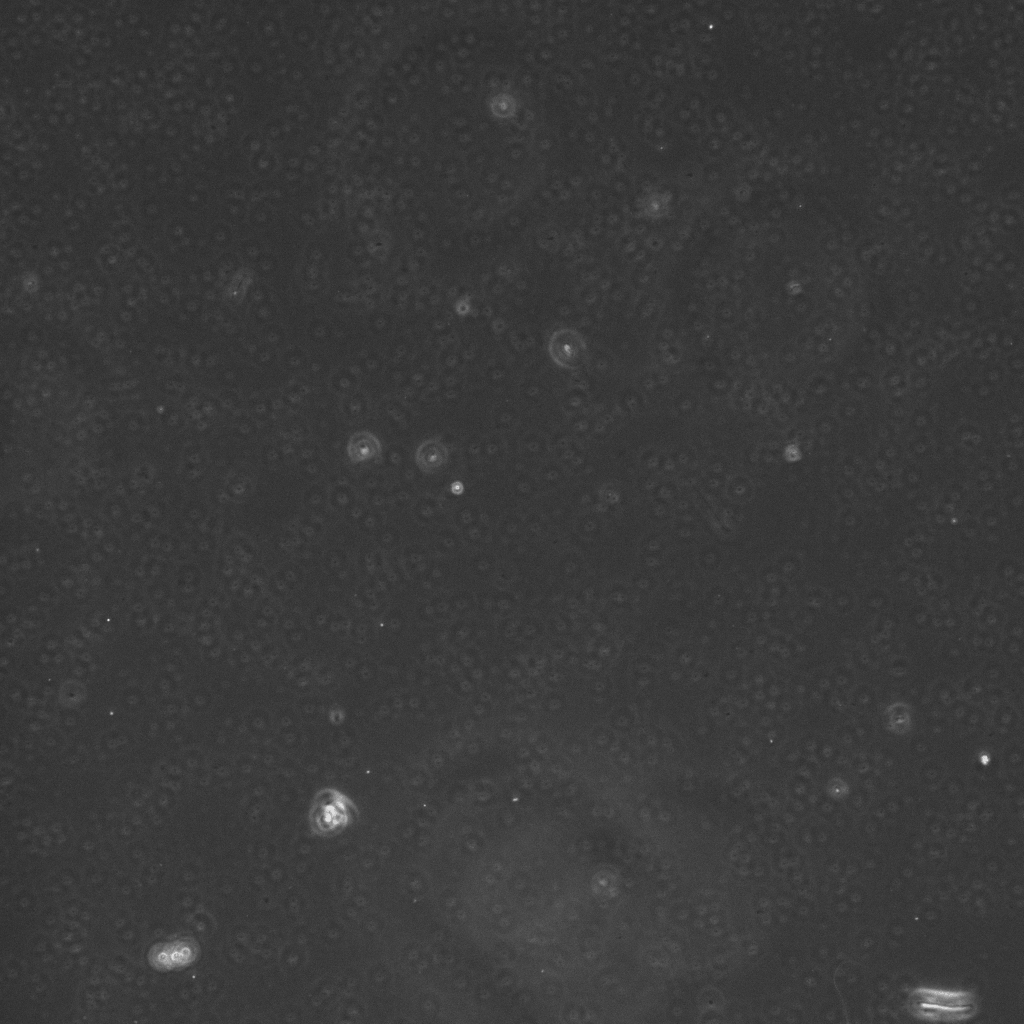

### Supplementary Video S3

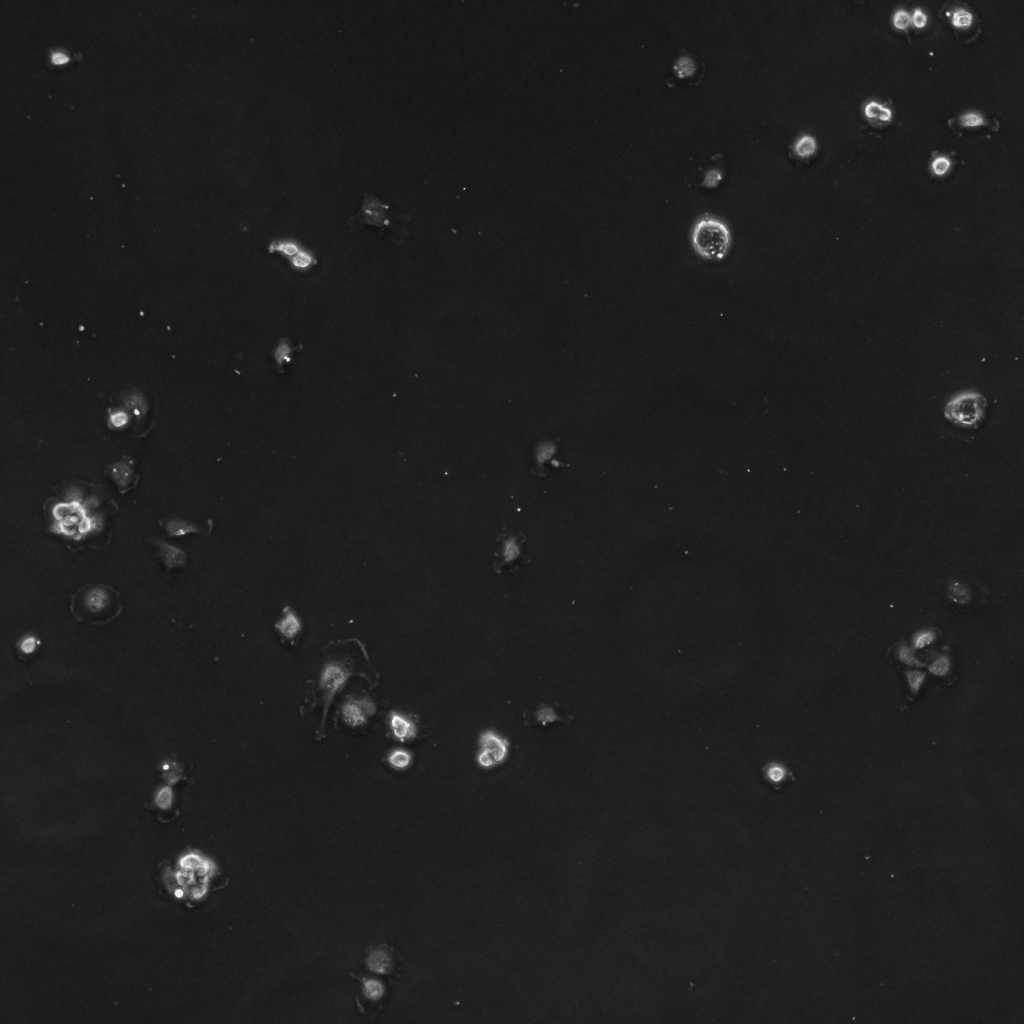
